# A Retina-inspired Optoelectronic Synapse Using Quantum Dots for Neuromorphic Photostimulation of Neurons

**DOI:** 10.1101/2023.09.30.560306

**Authors:** Ridvan Balamur, Guncem Ozgun Eren, Humeyra Nur Kaleli, Onuralp Karatum, Lokman Kaya, Murat Hasanreisoglu, Sedat Nizamoglu

## Abstract

Neuromorphic electronics, inspired by the functions of neurons, have the potential to enable biomimetic communication with cells. Such systems require operation in aqueous environments, generation of sufficient levels of ionic currents for neurostimulation, and plasticity. However, their implementation requires a combination of separate devices, such as sensors, organic synaptic transistors, and stimulation electrodes. Here, we present a compact neuromorphic synapse that combines photodetection, memory, and neurostimulation functionalities all-in-one. The artificial photoreception is facilitated by a photovoltaic device based on cell-interfacing InP/ZnS quantum dots, which induces photo-faradaic charge-transfer mediated plasticity. The device sends excitatory post-synaptic currents exhibiting paired-pulse facilitation and post-tetanic potentiation to the hippocampal neurons via the biohybrid synapse. The electrophysiological recordings indicate modulation of the probability of action potential firing due to biomimetic temporal summation of excitatory post-synaptic currents. Our results pave the way for the development of novel bioinspired neuroprosthetics and soft robotics and highlight the potential of quantum dots for achieving versatile neuromorphic functionality in aqueous environments.

## Introduction

Neurons are the building blocks of the nervous system and are responsible for generating, receiving, and transmitting electrical and chemical signals. They work in a fluid environment, communicate with synaptic transmission, and have plasticity via changing the strength of their response. By replicating the functionality of biological neurons, it is possible to achieve response capabilities that mimic human abilities [1]. Such biomimetic signal generation and transmission at bio-electronic interfaces can restore, replace, or enhance biological functions [2–5]. In addition, it can provide efficient event-driven processing and high-accuracy perception without separate memory and processing units [6, 7].

So far, neuromorphic conversion of exogenous signals (e.g., sound, pressure, light) into stimulation of neurons is achieved by bio-hybrid systems based on organic synaptic transistors, which advantageously offer biomimetic signal generation without bulky and rigid external encoding units [5]. These systems were used for tactile perception, vision-haptic fusion, and restoration of motion [3, 8-10]. They convert the external signals via an electronic sensor to electrical signals. Then, these electrical signals are applied to organic synaptic transistors for biomimetic plasticity. Finally, the output signals are applied to electrodes to stimulate neural tissues. However, such systems have been made of serial connections of separate elements such as sensors, transistors, and stimulation electrodes.

In this study, we introduce a novel biohybrid neuromorphic device, a quantum dot-based artificial synapse (QDAS), that combines photodetection, memory, and neurostimulation functionalities in a single compact device. QDAS employs InP/ZnS core/shell quantum dots (QDs), which directly interface with neurons and form a biohybrid synapse. The photovoltaic device elicits excitatory post-synaptic currents based on the Faradaic charge-transfer mechanism facilitating short-term plasticity under illumination. This plastic photoresponse emulates the temporal summation of biological neurons and enables modulation of neuron firing with memory. These results demonstrate that synaptic biointerfaces based on nanomaterials can enable unconventional forms of neuromorphic devices for future neuroprosthetics and soft robotics.

## Results

### Design of the quantum dot based artificial synapse (QDAS)

The concept of QDAS uses the direct conversion of light energy to neuromorphic ionic signals in the cellular environment for communication with neurons. All the light-to-ionic current conversion processes and neuromorphic signal generation are done by a single QDAS unit, which is analogous to the photoreceptor cells in the retina (Figure 1a). Biological photoreceptors absorb light and use neurotransmitters (i.e., glutamate) to convey information to secondary neurons (i.e., bipolar cells and horizontal cells). Secondary neurons communicate with the ganglion cells, which send action potentials to the brain via the optic nerve. In QDAS, light is absorbed by the InP/ZnS core/shell quantum dots, and the photogenerated electrons are transferred to the ITO via TiO_2_ layer while the hole is kept in the quantum dot (Figure 1b). In the device structure, TiO_2_ is crucial for enhanced exciton dissociation and effective electron transport (Figure 1c) [11]. The photovoltage on the quantum dots leads to a resistive (Faradaic) ionic photocurrent in the electrolyte, which is the postsynaptic current that induces action potentials by hippocampal neurons. Because of the Faradaic photoresponse, the neuromorphic signaling is triggered by the quantum dot and cell interface without any additional electrode. The photoactive side of the device acts as the stimulation electrode, and the ITO side acts as the return electrode that completes the current loop of the device-electrolyte circuit (Figure 1b). The flow of electric current between the QDAS and the cell leads to neurostimulation (Figure 1d). Hence, the neuromorphic device is effectively coupled with the primary hippocampal neurons.

**Figure 1.**
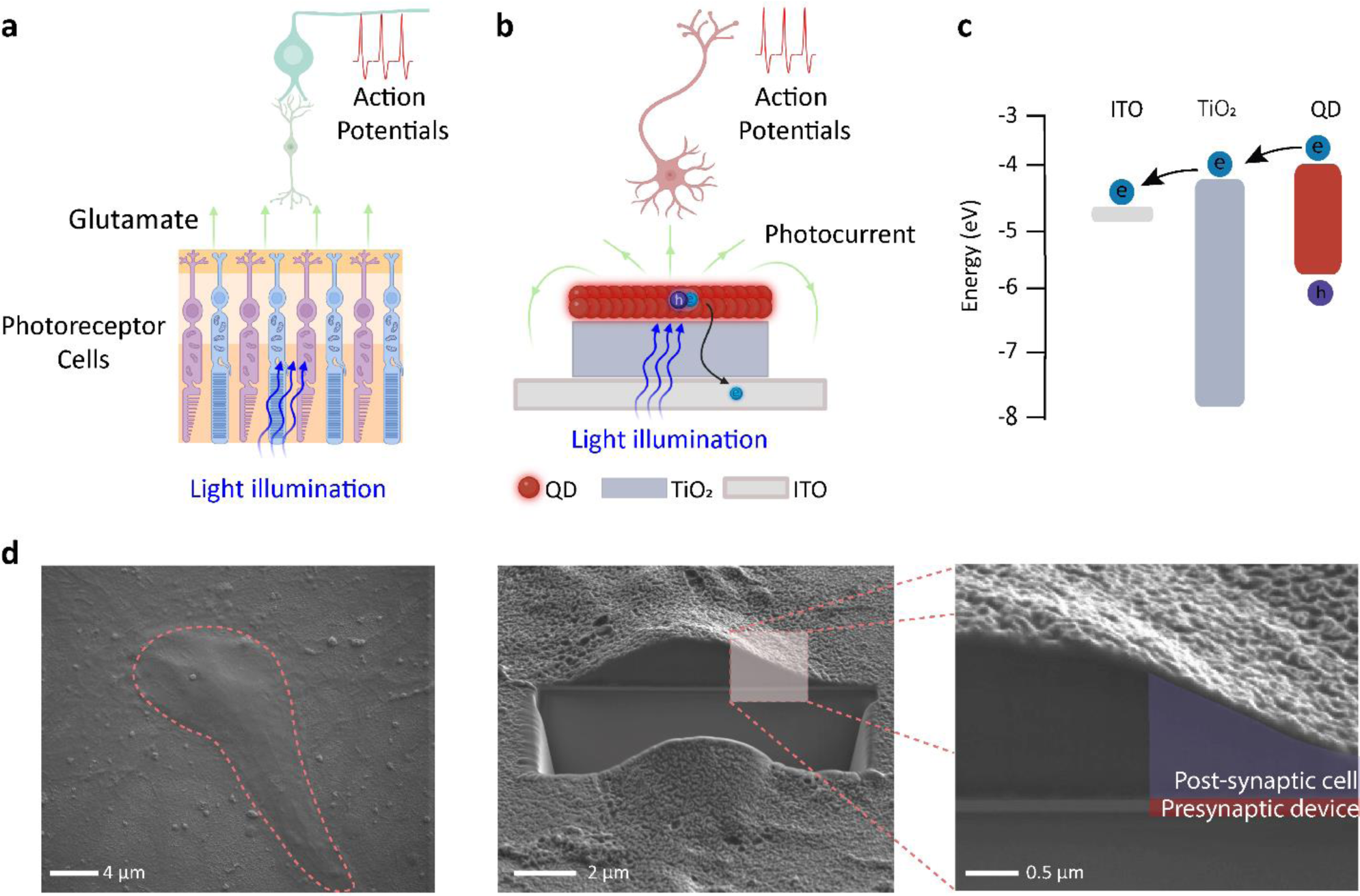
QDAS design and properties. **(A)** Schematic comparison of working principle of photoreception by the retina and **(B)** QDAS coupled with the hippocampal neurons. **(C)** Energy band diagram of QDAS [49, 50]. **(D)** SEM image of the (left) top view of the biointerface with the cultured hippocampal cells, (middle) FIB sectioned cell on the biointerface, and (right) QDAS (pre-synaptic device) coupled with the hippocampal neuron (post-synaptic cell).

### Optimization photoresponse of QDAS

To emulate the excitatory post-synaptic current (EPSC) response of biological neurons, the lifetime of photocurrents needs to be at least in the milliseconds range [12]. Since the decay time of capacitive current due to double-layer capacitance is fast in sub-millisecond range [13, 14], Faradaic charge transfer is a suitable option that allows injection of ionic currents during the light turn-on periods and decays slowly in milliseconds range. In comparison with AlSb QDs generating capacitive currents [15], InP QDs are appropriate materials due to the Faradaic currents with longer lifetime [16–18] and the three-electrode electrochemical characterization setup (in Figure 2a) is utilized to measure photocurrent characteristic of the QDAS.

**Figure 2.**
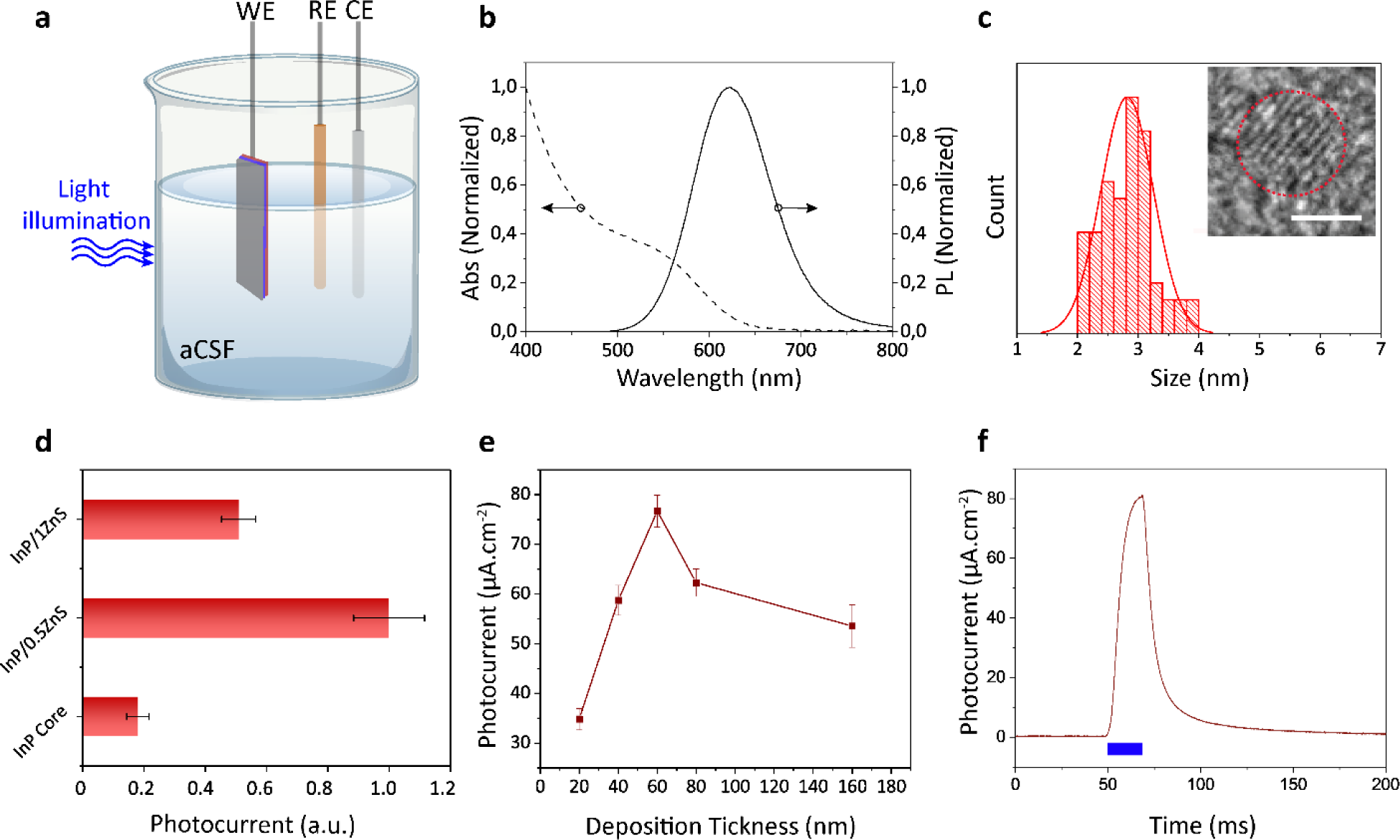
Photoelectrochemical characterization, optimization, and measurements. **(A)** Schematic of photo-electrochemical setup. The setup included QDAS as the working electrode, a platinum counter electrode, and an Ag/AgCl reference electrode. Illumination was applied to the 1 cm^2^ area of the device. **(B)** Absorption and PL spectrum of InP/0.5ZnS core/shell QD. **(C)** Size distribution of InP/0.5ZnS core/shell QDs. The inset shows the HR-TEM of InP/0.5ZnS. (Scale bar: 2 nm). 200 QDs are counted for size distribution. **(D)** Photocurrent response of InP core, InP/0.5ZnS, and InP/1ZnS core/shell QDs. **(E)** Photocurrent response of QDAS with InP/0.5ZnS core/shell QDs for different QD layer thicknesses. The photocurrent is measured under 450 nm light pulses with 20 ms pulse duration (means ± SD, n=5). **(F)** Photocurrent response of optimized QDAS under 20 ms, 450 nm light illumination.

InP core quantum dots (QDs) were synthesized using an amine-derived synthetic approach, with a mean size of 2.49 nm [19]. A 0.16 nm thick ZnS shell, corresponding to 0.51 monolayers, was grown on top of the InP core using successive ionic layer adsorption and reaction (SILAR) method to investigate the effect of the shell on the photocurrent [20]. The InP/0.5ZnS QDs have a cubic crystal structure (Figure S1), and their photoluminescence is in the red spectral region (Figure 2b, c). The shell growth reduces trap states, which is evident from the increase in the photoluminescence quantum yield (PLQY) from 10.7% to 16.1%. Compared to InP core QDs, InP/0.5ZnS core/shell QDs exhibit a significant enhancement in photocurrent peak (Figure 2d). However, when the ZnS shell thickness is further increased to 0.33 nm (Figure S2), corresponding to 1.05 monolayers (i.e., InP/1ZnS), the photocurrent peak starts to drop. This is due to the decrease in the tunneling probability of photogenerated charges by the thicker potential barrier of ZnS. Hence, we select to use InP/0.5ZnS core/shell QDs for QDAS.

To achieve reduced interparticle spacing for high conductivity and colloidal stability of the QD film in aqueous environments, the large native ligands of hexadecylamine (HDA) are replaced with the short-chain ligand of 3-mercaptopropionic acid (MPA) via solid-state ligand exchange. After QDs are coated on the TiO_2_, the process involves using spin-coating of QDs, soaking to the solution with MPA for ligand exchange and washing with protic solvent to remove excess new short ligands. MPA also functions as a hole acceptor on the QD surface, which is beneficial for the photovoltaic effect that can induce potential difference between working and return electrodes for photocurrent generation [21]. Different QD layer thicknesses are investigated to understand the photocurrent levels in artificial cerebrospinal fluid, which is close to the condition of the cellular environments (Figure 2e). If the photoactive layer is thick, extracted charges recombine in the neutral region, which decreases the charge extraction efficiency. On the other hand, thin photoactive layer is disadvantageous due to low absorption. We obtained the maximum photocurrent when the QDs layer thickness is 60 nm. A single spike of light illumination can trigger an EPSC of 80 µA.cm^-2^ (Figure 2f), and EPSC decays to the baseline level within tens of milliseconds because of the surface-voltage drop after recombination and discharge through the return electrode.

To understand the source of photocurrent, we did electrochemical analysis of QDAS in aCSF solution by removing the ingredients one by one. We observed that when HEPES is removed, the photocurrent significantly decrease, and this shows that HEPES oxidation is an essential enabler for Faradaic photocurrent generation (Figure S3) [22]. As a control, we removed QDAS, and when we measured the photocurrent of only TiO_2_-coated ITO devices, no detectable photocurrent was observed, indicating that the HEPES photooxidation process was triggered by QDAS. Therefore, water [23] and HEPES oxidations by QDAS are possible sources of electrochemical photocurrent.

### Synaptic characteristics of QDAS

In biological neurons, the action potentials of presynapse trigger the release of the neurotransmitters across the synaptic cleft and lead to the generation of action potential in the post-synapse with plasticity. Likewise, the light pulses induce the accumulation of holes on the device surface, and the positive photovoltage by the neuromorphic device leads to a postsynaptic Faradaic current for neurostimulation with short-term memory. When subsequent inputs are applied to the QDAS, the charges cannot find sufficient time to discharge across the return electrode, and the amplification of the EPSC is observed (Figure 3a). The amplitude of EPSC from the second spike (A_2_) is almost doubled with respect to the first spike (A_1_), and the EPSC saturates to a level of 40 µA.cm^-2^ after eight spikes. This accumulation of EPSC signals can be modulated by changing the illumination frequency. When the illumination frequency increases from 5 to 100 Hz, the amplification of EPSC signals increases from 19 µA.cm^-2^ to 100 µA.cm^-2^, respectively (Figure 3b). Each applied pulse for different frequencies has 5 ms on time and frequency depends only on off-time of the illumination. Therefore, the saturation level depends on the duration of the off-time, where longer durations will cause more relaxation of photocurrent, and cause lower saturation current for that specific frequency. Such amplified signal generation is also observed by biological neurons in the hippocampus, cortex, and spinal cord [24, 25].

**Figure 3.**
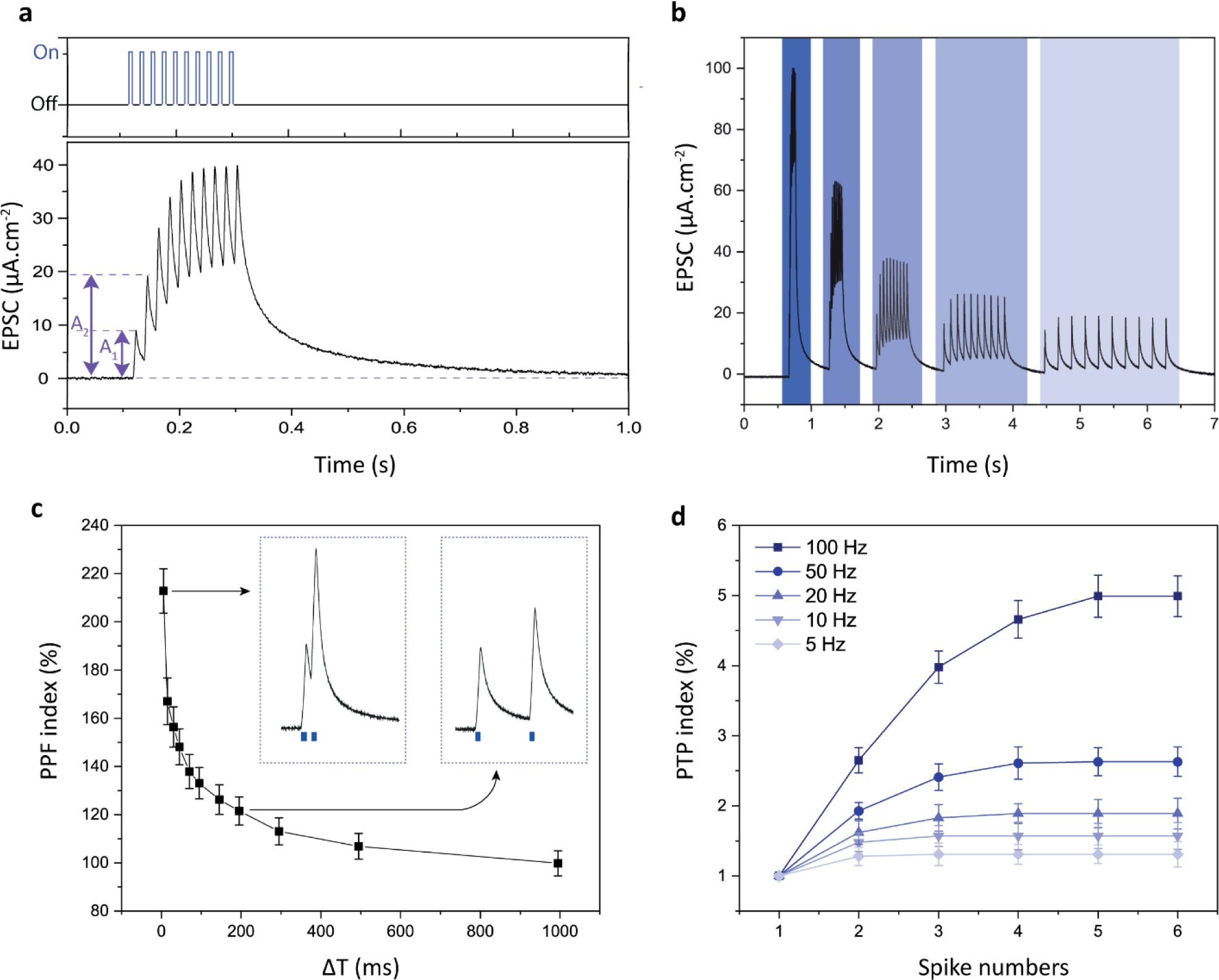
Photo synaptic characterization of QDAS. **(A)** (Top) The applied illumination pulse train and (bottom) the resulting EPSC. **(B)** EPSC under light pulse train consisting of ten spikes with frequencies of 100, 50, 20, 10, and 5 Hz (from left to right) (off time: 5, 15, 45, 95, 195 ms, respectively, and on time: 5 ms). **(C)** PPF index (%) is defined as 100% × A_2_/A_1_ for different off-times (ΔT). The on-time is 5 ms. The left inset shows EPSC with ΔT=5 ms, and the right inset shows ΔT=195 ms. **(D)** PTP index (%), defined as 100% × A_n_/A_1_, is plotted as a function of the number of spikes (n = 1:6) for different frequencies. A_n_ and A_1_ represent the amplitudes of the n^th^ and first pulses, respectively. The on-time is 5 ms.

The PPF index measures the short-term plasticity depending on the applied input, and similar to biological neurons, QDAS displays stronger PPF index when two consecutive spikes are applied with smaller off times (Figure 3c). For example, while the pulses with the off-time of 195 ms are applied, the accumulation of photocurrent leads to a PPF of 121%, and if the off-time decreases to 5 ms, PPF increases to 213%. The PTP index was also investigated for different frequencies (Figure 3d), and the PTP index increases with the higher frequency illumination trains. For example, while the PTP index is 131% at 5 Hz illumination, it can reach up to ∼500% at 100 Hz. High-frequency action potential firing elevates the synaptic strength of biological neurons, and analogously QDAS also increases the photocurrent response in high-frequency bursts of light illumination.

### Biocompatibility

Biocompatibility is an important need for biointerfacing with neurons, and the composition of cell interfacing QDs directly affects biocompatibility. Brain cells, especially in the hippocampal region, are cell populations widely used in electrophysiology studies and allow the evaluation of action potential abilities in different conditions. If the devices would induce toxicity on cells, the stress on neurons would be decremental on electrophysiology. Herein, we used hippocampal neurons isolated from rat embryos at embryonic days 15-17 for cell culture and patch clamp experiments. The neurons were cultured on QDAS and ITO control substrates for viability and morphology analyses to examine the short-term effect after 3-days of incubation. On 3-days of cell culture, a slight reduction of cell viability was observed on QDAS substrates compared to the control substrates (Figure S5). In terms of inorganic nanostructure, the ZnS shell has non-toxic elements, leading to even high biocompatibility for cadmium-based quantum dots [26, 27]. Here we consider that the slight decrease of biocompatibility may be due to TiO_2_ [28–30]. Cytoskeleton and adult neuron marker stainings were performed on day 14 of culture to check long-term cell characteristics and morphology changes. Cells were fixed after 3-day and 14-day cultures and stained with NeuN and f-actin filaments to visualize nucleus, cytoskeletal and network connections. According to the results, cells can establish enhanced neurite connections on neuromorphic devices as on control substrates and adapt to the long-term culture condition (Figure S6). Before performing electrophysiology experiments, we checked the stability of the device after 14 days in culture environment (Figure S8) and observed sufficiently high-level of photocurrents, 80% of the initial photocurrent level, for neurostimulation.

### Biohybrid neurostimulation

For biohybridization, we cultured primary hippocampal neurons on the synaptic device. To investigate the coupling of the neuromorphic device with biological neurons, we used patch clamp electrophysiology to record the single-cell electrical activity under illumination (Figure 4a). The intracellular potential of the cultured cell is measured from the cell soma (Figure 4a inset) with respect to a distant Ag/AgCl bath electrode in the whole-cell configuration without applying any bias current during the measurements. The cells show a mean resting membrane potential of −65 mV, and cells can go into a non-linear response as an action potential (AP) with a depolarization of 25 mV (Figure 4b).

**Figure 4.**
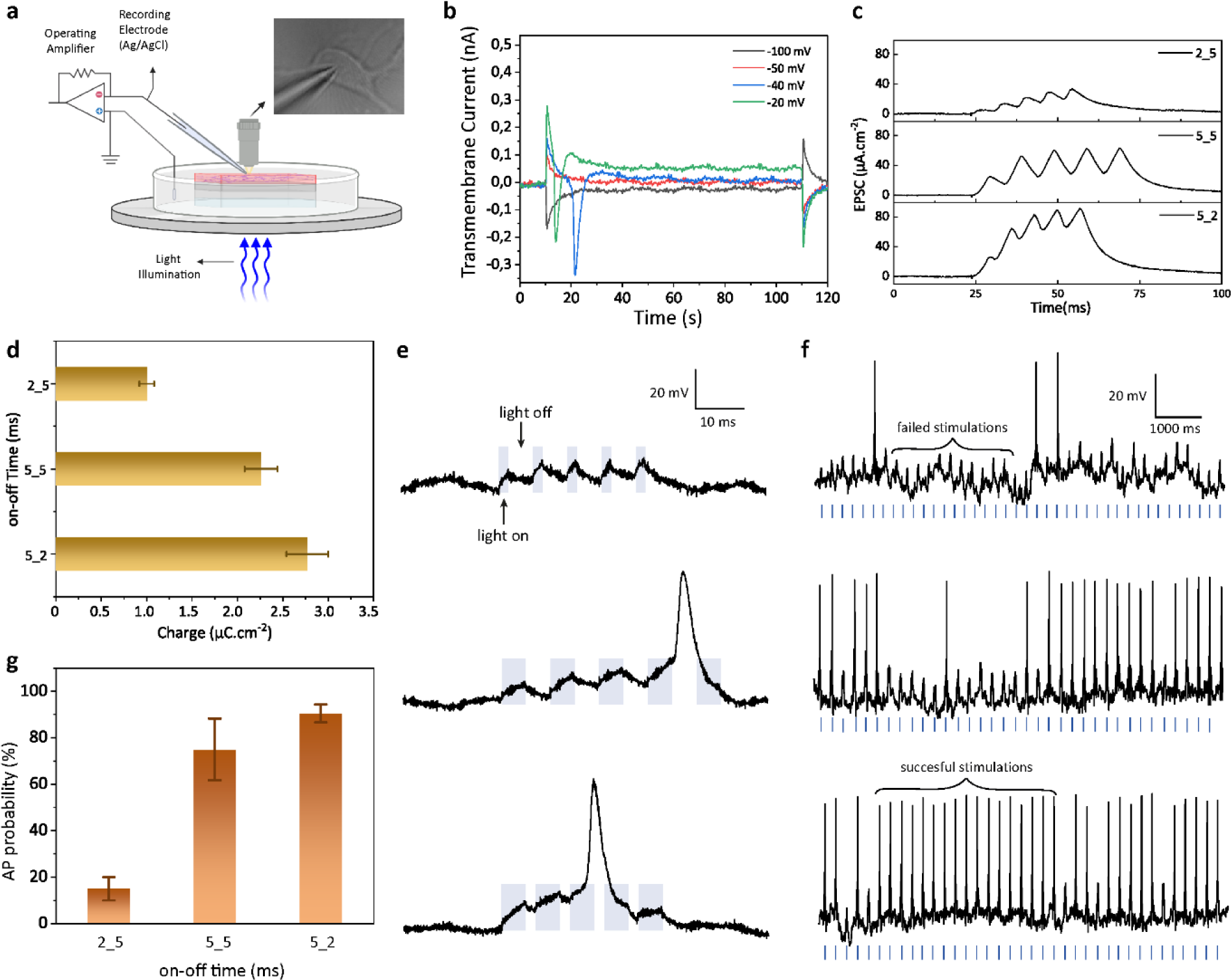
Single cell electrophysiology recordings of QDAS. **(A)** Schematic of the patch-clamp electrophysiology recording setup. The inset shows a photograph of an exemplary patched hippocampal neuron. **(B)** Recorded transmembrane currents in whole-cell configuration for different holding potentials. **(C)** Probability of generating action potential (AP) for different on and off cycles (mean ± SEM, n=3). **(D)** EPSC signal triggered with different illuminations, (top) illuminated with 2 ms on-time and 5 ms off-time, (middle) illuminated with 5 ms on-time and 5 ms off-time, (bottom) illuminated with 5 ms on-time and 2 ms off-time. **(E)** Photogenerated charge with different illumination on-off cycles calculated from (d). **(F)** Membrane potential under a single light signal. The signal consists of five successive pulses with varying on- and off-times (x_y). **(G)** Membrane potential under the light train of the signals in (F) at a frequency of 5 Hz. The top, middle, and bottom panels correspond to x_y values of 2_5, 5_5, and 5_2, respectively, where x and y denote the on- and off-times in milliseconds.

To show the coupling of synaptic plasticity to the biological neurons, we use light signal configurations in milliseconds time scale, compatible with the electrophysiological activity of neurons. Each light signal configuration is composed of five successive light pulses with varying on- and off-times (i.e., x_y where x is the on-time and y is the off-time in milliseconds) and transduces clearly observable EPSC peaks for each light turn-on period (Figure 4c). When QDAS is repeatedly stimulated within a short time period, the inputs add up over time, mimicking temporal summation by biological neurons (Figure 4d) [31]. The summation was achieved by flashing light on the neuron with a repetition period shorter than the decay time, which is not possible via currents due to double-layer capacitance (Figure S9). While the condition of light pulse configuration changes from 2_5 to 5_5 and 5_2, the peaks and the total charges generated by the EPSC increase because of the biomimetic response.

Since the surface charge of the QDAS is positive under illumination due to the confinement of the photogenerated holes, QDAS induces EPSC from the QD surface toward the electrolyte that induces hyperpolarization of the attached membrane and depolarization of the recorded free membrane [32] (Figure 4e and 4f). Like in EPSC, the depolarization peaks of the membrane potential are also observable after each flash of light (Figure 4e). For the light signal configuration of 2_5, the membrane can only depolarize 13-mV after five successive light pulses (Figure 4e, top)) and thus, the probability of obtaining action potential is low, which is 15% (Figure 4g and 4f, top)). Increasing the on-time to 5 ms while keeping the off-time fixed at 5 ms (i.e., 5_5) strengthened the EPSC signal (Figure 4c, middle) and increased the depolarization in the membrane potential (Figure 4e, middle). Thus, the probability of generating an action potential significantly increased to 75% (Figure 4f, middle and 4g)). To further boost the EPSC signal, the off-time is reduced to 2 ms while maintaining the on-time constant at 5 ms (i.e., 5_2) (Figure 4c bottom), and this pulse configuration was able to produce action potentials with a probability of 90% (Figure 4e and 4f bottom, 4g). A glass/ITO substrate was employed as a control device, and there was no depolarization or action potential production with the light irradiation on the control device. Moreover, the light intensity level (30 mW.cm^-2^) is advantageously below the ocular safety limits [33].

### Discussions and conclusion

The QDAS’s compactness and wireless operation make it advantageous for implantation into vulnerable tissues. The use of wires and interconnected receiving and processing electronics during surgery can be a significant source of difficulty and may lead to unwanted side effects [34]. Previous studies have shown that photovoltaic retinal implants can be implanted into the sub-retina [35] and epi-retina [36], demonstrating the potential of photovoltaic devices for implantation. Compared to wired systems such as Argus II and Alpha AMS, which require 4 hours of surgery time, PRIMA without any wires can be implanted in 1-2 hours [37]. Hence, compact, and wireless devices hold high promise for future neuroprosthetics.

Sufficiently high level of charges is generated for neurostimulation because of the Faradaic nature of the current. Even though the capacitive currents are the preferred way of stimulating neurons for clinical application, electrochemical currents have been utilized for neuronal excitation in previous studies [14, 38, 39] and we also observed precise and repetitive firing of neurons. The transparency of the eye is advantageous for generating sufficiently strong photocurrent levels in the retina by direct illumination of external light sources. In addition, injection of the achievable charge levels by QDAS in the range of tens of µC.cm^-2^ can facilitate stimulation of a wide variety of tissues, including the sciatic nerve, subthalamic nucleus, and cortical neurons [39–41]. Moreover, biocompatible optical fibers can be used to deliver light to the QDAS when it is implanted into non-transparent parts of the body [42–44].

Biomimetic neurostimulation devices can be combined with artificial sensing [45] and spiking neurons [46, 47] to develop advanced neuromorphic systems. Previously, electrochemical transistors were used to sense neurotransmitters such as dopamine and regulate synaptic plasticity through channel conductance variation [45]. These transistors also enabled the development of neuron-like spiking circuits controlled by ion concentration [5, 45]. Combining ionic sensing, spiking, and receiving actuation signals (e.g., light, sound, and motion) with neurostimulation can lead to organ-like closed-loop neuromorphic systems. This strategy can be beneficially used for future neuroprosthetics and miniaturized robots to rehabilitate neurological disorders.

In conclusion, we showed that cell-interfacing quantum dots enabled photon detection, synaptic and neurostimulation functionalities in one single compact device architecture. The device imitated the short-term plasticity of biological neurons via Faradaic currents and converted light pulses to neuromorphic photostimulation of neurons. Hence, nanomaterials show high promise as a building block for unconventional and biomimetic control of cells.

## Materials and Methods

### Experimental Section

Indium chloride (InCl_3_) (99.99%, trace metals basis), hexadecylamine (HDA-98%), zinc undecylenate (98%), 1-octadecene (ODE-technical grade, 90%), stearic acid (SA) (98%), tris(trimethylsilyl)phosphine (P(TMS)_3_) (95%), sulfur (99.98%, trace metals basis), zinc stearate (Zn(St)_2_-technical grade), and trioctylphosphine (TOP-97%) were used from Sigma Aldrich.

### P(TMS)_3_ stock solution preparation

A solution of P(TMS)_3_ in ODE was prepared with the concentration of 0.1 mmol.ml^-1^ and stored in inert atmosphere at −30 °C.

### Zinc (Zn) and sulfur (S) stock solutions preparation

For Zn precursor solution 558 mg zinc stearate (Zn(St)_2_) was dissolved in 8 mL ODE; for the S precursor solution 32 mg sulfur was dissolved in 8 mL TOP.

### Synthesis of InP Core QDs

For the synthesis of InP core QDs, 22 mg (0.1 mmol) InCl_3_, 48 mg (0.2 mmol) hexadecylamine (HDA), 43 mg (0.1 mmol) zinc undecylenate and 28 mg (0.1 mmol) stearic acid (SA) were mixed in 3 ml ODE in a three-neck 50 ml round bottom reaction flask. The solution was firstly degassed at 120 °C for 1.5 h, then heated up to 240°C under nitrogen atmosphere. Afterwards, 0.5 ml of the P(TMS)_3_ from stock solution (0.1 mmol) was injected to the main solution at this temperature. The solution was kept at 230°C for 20 min and then cooled down to room temperature. InP core solution was centrifuged at 9000 rpm, 20 min using hexane-ethanol. Centrifugation process was repeated 2 times. The final InP core QD solution was redispersed in toluene.

### Synthesis of InP/ZnS core/shell QDs

To synthesize InP/ZnS core/shell QDs with multiple ZnS shell, InP core solution was cooled down to 150°C. For ZnS shell growth, zinc and sulfur precursor solutions (in Table S1) were added, and the solution temperature was increased to 220°C. The solution was mixed at 220°C for 40 min. For additional shells, same methodology was performed. Finally, InP/ZnS core/shell solution was cooled down to room temperature and the resulted solution was centrifuged at 11000 rpm, 3 min. To minimize the amount of zinc-based organic parts, centrifugation at 11000 rpm was repeated 3 times [48]. Then, the QD solution was mixed with hexane and ethanol with a volume ratio of 1:2:4 and precipitated at 9000 rpm for 10 min. Precipitation process was repeated 2 times. The final QDs solution was redispersed in toluene.

### Structural and optical characterizations of QDs

Powder X-ray diffraction (XRD) of InP/ZnS core/shell QDs were carried out using a Rigaku Miniflex XRD λ_Cu Kα_ = 1.54 Å with a 5° min^−1^ scan rate. XRD measurements were carried out at room temperature. XRD samples were prepared by drop-casting QDs dissolved in hexane onto 1 cm × 1 cm silicon wafers and heated to 100°C and kept at this temperature for 1 h to dry liquid QDs on the silicon wafers. DF-STEM images were collected using Hitachi HF5000 TEM/STEM operated at 200kV. For TEM measurement, samples were deposited as 10 μL of a 1 mM QD in a hexane solution on a copper support grid.

An Edinburgh Instruments spectrofluorometer FS5 with a 150 W xenon lamp was used to measure absorbance and PL. The excitation wavelength was 375 nm. For absorbance and PL measurements, regular quartz cuvettes were used. PLQY measurements were carried out using an integrating sphere module with an inner diameter of 150 mm. The integrating sphere was inserted into the FS5 device for the measurement of quantum yields.

### Biointerface fabrication

Biointerfaces were fabricated on glass substrates, which are coated with indium tin oxide (ITO) (Sigma-Aldrich 639303). Four step cleaning procedure of substrates starts with sonication in deionized water solution with 3% Hellmanex-3, and continuous with deionized water, pure ethanol, and isopropyl alcohol each for 15 min at 50°C. To dry the cleaned substrates, they were kept in oven for 20 min at 40°C. Before coating the TiO_2_ layer, cleaned substrates were treated with UV-ozone for 25 min. TiO_2_ paste (Sigma-Aldrich) was diluted with ethanol 1:9 mass ratio and sonicated for 4 hours. Then, to obtain a more homogenous TiO_2_ layer, 2 layer was coated by spin coating in 2000 rpm with 20 mins of 50°C annealing in between. After the second layer the substrates were annealed at 50°C for 5 mins, 100°C for 5 mins, 200°C for 5 mins, and at 400°C for 45 mins. QDs layer was coated on TiO_2_ with layer-by-layer deposition. Each QD layer coating consists of 3 steps; first 50 mg.ml^-1^ QDs in toluene were spined at 1000 rpm, then 1M 3-MPA methanol solution was coated with at 1000 rpm, the surface was washed with pure methanol by spin coating at 1000 rpm. After the desired thickness is obtained, the devices were annealed at 110°C for 10 mins. The thickness of the QD layers is characterized using cross-sectional SEM, Zeiss ultra plus field emission scanning microscope.

### Photo-electrochemical measurements

Autolab Potentiostat Galvanostat PGSTAT (Metrhom, Netherlands) was used for the photocurrent and photovoltage response of our device. The three-electrode configuration utilizes of Ag/AgCl as the reference electrode, platinum wire as the counter electrode and connection to the ITO layer of the biointerfaces as the working electrode inside the extracellular medium (artificial cerebrospinal fluid (aCSF)). The aCSF medium was prepared by mixing 10 mM 4-(2-hydroxyethyl)-1-piperazineethanesulfonic acid (HEPES), 10 mM glucose, 2 mM CaCl_2_, 140 mM NaCl, 1 mM MgCl_2_, 3 mM KCl, and a stoichiometric amount of NaOH to adjust the pH to 7.4, in distilled water. Exactly 1 cm^2^ of the substrate was immersed into aCSF solution to measure the current density. The light pulses were applied with Thorlabs M450LP1 LED. The optical power was controlled with an optical power meter (Newport 843-R). The optical power density of the applied light illumination in Figure 2 and 3 was 30 mW.cm^-2^. The data was analyzed using the NOVA software.

### Primary Neuron Isolation

All experimental procedures have been approved by the Institutional Animal Care and Use Committees of Koç University (Approval No: 2021.HADYEK.022) according to Directive 2010/63/EU of the European Parliament and of the Council on the Protection of Animals Used for Scientific Purposes. For isolation of primary hippocampal neurons, brains of Wistar Albino rats at embryonic day 15-17 (E15-E17) were dissected and blood vessel layer was removed carefully from brain. Cortexes were divided and hippocampi were dissected. Tissues were placed in ice-cold Hank’s Balanced Salt Solution (HBSS, Thermo Fisher Scientific, MA, USA) supplemented with penicillin/streptomycin, glucose, and sucrose. After dissection, samples were washed twice with ice cold HBSS and they were incubated in 0.25% Trypsin-EDTA solution (Thermo Fisher Scientific, MA, USA) supplemented with 2% DNase-I (NeoFroxx, Einhausen, Germany) for enzymatic digestion for 20 minutes in a 37°C water bath. After incubation, Dulbecco’s Modified Eagle Medium/Nutrient Mixture F-12 (DMEM/F12 Thermo Fisher Scientific, MA, USA) supplemented with 10% fetal bovine serum (FBS, Heat Inactivated, GE Healthcare, IL, USA) and 1% penicillin/streptomycin (Thermo Fisher Scientific, MA, USA) was added to inhibit excess enzyme activity and tissue suspension was centrifuged for 3 min at 300 rcf. Supernatant was removed and plating Neurobasal Medium (NBM, Thermo Fisher Scientific, MA, USA) supplemented with B27, L-glutamine, penicillin/streptomycin, β-mercaptoethanol and glutamate was added on tissues. For complete digestion, tissues were triturate with a plastic Pasteur pipet for 10 times. Digested tissue suspension was passed through a 70 µm cell strainer. The homogenous cell solution was collected in new falcon tube, and they were cultured on poly-l-lysine (PLL, Sigma-Aldrich, MO, USA) coated ITO and Neuromorphic biointerfaces. Cells were incubated for 3 days at 37°C incubator with 5% CO_2_. After 3-day incubation, media was refreshed with NBM supplemented with cytosine arabinoside (Sigma-Aldrich, MO, USA) to eliminate glia proliferation. After 1-day incubation with glia inhibition media, fresh NBM was replaced to continue culturing for biocompatibility and electrophysiology experiments. Half of media were changed every 3-4 days.

### Biocompatibility Assay

Cells’ viability cultured on control ITO and NM devices was examined with CellTiter-Glo Viability Assay (CTG, Promega, Mannheim, Germany). After sterilization of the devices with 70% ethanol followed by UV irradiation for 30 min, substrates were placed in the 6-well plates and they were coated with poly-l-lysine (PLL, Sigma-Aldrich, MO, USA) solution prepared in ddH2O. Isolated primary hippocampal neurons were seeded as 500000 cells/well on PLL coated devices. After 3 days of incubation, AraC glia removal medium was added for 24 hours. After glia inhibition, fresh NBM was replaced to maintain culturing at 37 °C and 5% CO_2_. Next day 1:1 ratio of CTG solution was added to cells cultured on substrates to check cell viability. Cells were incubated with CTG solution for 10 minutes at room temperature. The solution was transferred to 96-well opaque plate and the luminescence signals was measured with Synergy H1 Microplate Reader (Bio-Tek Instruments). The relative cell viability of each sample was calculated as follows: Viability = (ODsample/ODcontrol) × 100. The optical density (OD) of the sample was obtained from the cells grown on the Neuromorphic substrates, and the OD of the control was obtained from the cells grown on the ITO control substrates.

### Immunofluorescence Staining and Imaging

Primary hippocampal neurons were seeded as explained above on the ITO control and the Neuromorphic devices as 500000 cell/device. Short-term effect of samples on cell culture was checked at Day 0 and long-term alteration was monitored on Day 14. The neurons at Day 0 or Day 14 were fixed by cold 4% paraformaldehyde in PBS for 20 min and washed three times with PBS-T (Phosphate Buffered Saline, 0.1% Triton X-100). For permeabilization, the cells were incubated with 0.1% TritonX-100 in PBS for 8 min at room temperature. After permeabilization, cells were incubated with Superblock (Thermo Fisher Scientific, MA, USA) for 10 min at room temperature. For neuron characterization, mature neuron marker NeuN was used. For labelling of neurons, overnight incubation with anti-NeuN antibody (ab177487, Abcam, Cambridge, UK) in blocking solution was applied. After incubation, samples were washed with PBS-T and they were incubated with goat anti-rabbit IgG H&L Alexa Fluor 488 (Cell Signaling Technology, MA, USA) for 90 minutes at 37°C. To show morphology of neurons cultured substrates, they were labeled with FITC-conjugated phalloidin antibody for 90 minutes at 37°C. All samples were washed three times with PBS-T, then mounted with DAPI supplemented mounting medium (50001, Ibidi GmbH, Germany). All immunofluorescence images were taken by inverted fluorescence microscope (Axio Observer Z1, ZEISS, Oberkochen, Germany).

### Sample Preparation for FIB/SEM

Cells grown on QDAS were washes with ddH2O to remove cell debris and death cells. Cells were fixed with EM grade 2% glutaraldehyde and 2.5% paraformaldehyde prepared in 0.1 M Sorenson phosphate buffer for 2 h at room temperature. After fixation cells were washed with 0.1 M Sorenson phosphate buffer 3 × 10 min. Afterward, samples were dehydrated in ethanol series of 30%, 50%, 70%, 90%, 95%, 100% ethanol prepared with ddH2O for 10 min respectively. Samples were allowed for overnight air drying before SEM imaging. To remove charging of materials, the samples were sputter-coated with thin layer of gold before SEM imaging. Top view and cross-sectional SEM images of single neuron soma on QDAS were taken by the Hitachi Ethos NX5000 Focused Ion Beam (FIB) (Hitachi, Japan). Ultrahigh resolution was obtained at 3 kV. For cross sectional view and to check layers of QDAS, central area of cell was etched with FIB system.

### Electrophysiology Experiments

Experiments were performed by EPC 800 patch clamp amplifier (HEKA Electronic GmbH, Pfalz, Germany). QDAS integrated and control devices were cleaned with 70 vol% ethanol solution and incubated for 3 days in DI water. The pulled patch pipettes of 4–6 MΩ were utilized to carry out the whole-cell patch clamp experiment. The internal cellular medium was prepared by mixing 140mM KCl, 2mM MgCl_2_, 10mM HEPES, 10mM ethylene glycol-bis (β-aminoethyl ether)-N,N,N′,N′-tetraacetic acid (EGTA), 2mM Mg-ATP in water, and the pH was calibrated to 7.2–7.3 using 1M KOH. The intracellular solution was filled into the patch pipettes to achieve whole-cell patch. A digital camera integrated Olympus T2 upright microscope was used to patch and monitor the cells. The blue LED (M450LP1, Thorlabs Inc, NJ, USA) was used as the light source. LED system was driven by DC2200 - High-Power 1-Channel LED Driver (Thorlabs Inc., NJ, USA).

## Acknowledgements

We thank Dr. Baris Yagci from the Koc University Surface Science and Technology Center (KUYTAM) for the SEM images. We thank Dr. Amir Motallebzadeh from KUYTAM for helping with oxygen plasma sterilization. We thank Dr. Gulcan Corapcioglu for HR-TEM images. The authors gratefully acknowledge use of the services and facilities of the Koç University Research Center for Translational Medicine (KUTTAM), funded by the Republic of Turkey Ministry of Development. The content is solely the responsibility of the authors and does not necessarily represent the official views of the Ministry of Development.

## Author contributions

Conceptualization: SN, MH

Methodology: SE, MH, RB, GOE

Investigation: SE, MH, RB, HNK, LK

Visualization: RB, HNK

Supervision: SN, MH, OK

Writing—original draft: SN, RB, HNK

Writing—review & editing: SN, RB, HNK

---

**Table S1.** Zinc and sulfur precursor solutions used for ZnS shelling.

|  | **Concentration**<br>**(M)** | **InP/0.5ZnS** | **InP/1ZnS** |
| --- | --- | --- | --- |
| **ZnSt<sub>2</sub>-**<br>**ODE** | 0.1 | 600 μL | 1050 μL |
| **S-TOP** | 0.1 | 600 μL | 1050 μL |

**Figure S1.**
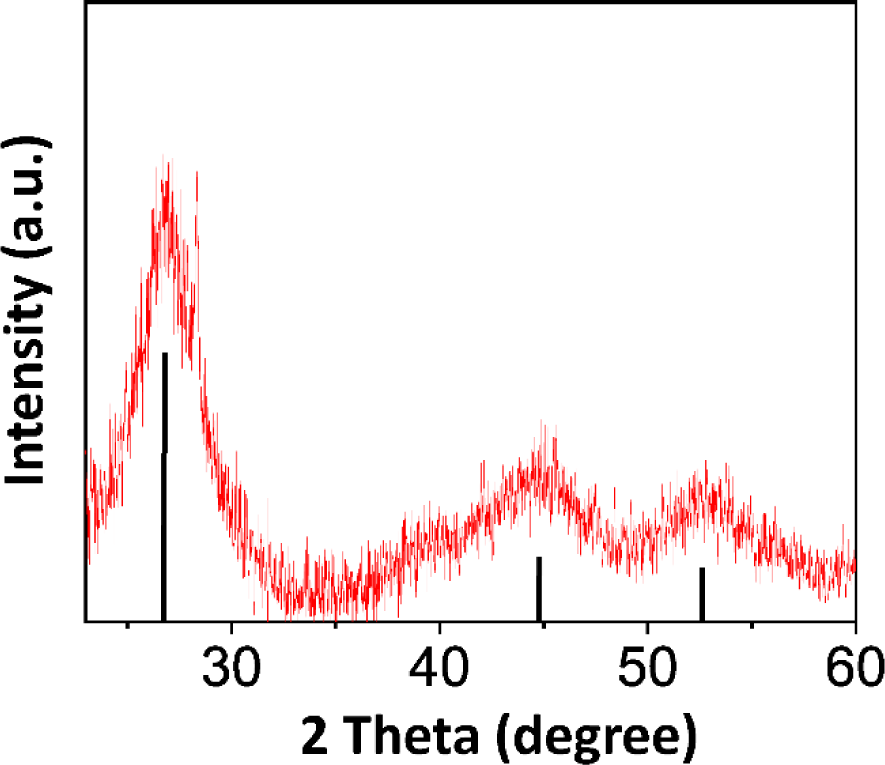
XRD pattern of the InP/ZnS core/shell QDs. The X-ray diffraction (XRD) pattern reveals the crystal planes of the (111), (220), and (222) of the QDs. (InP JCPDS No. 32-0452 and ZnS JCPDS No. 80-0020)

**Figure S2.**
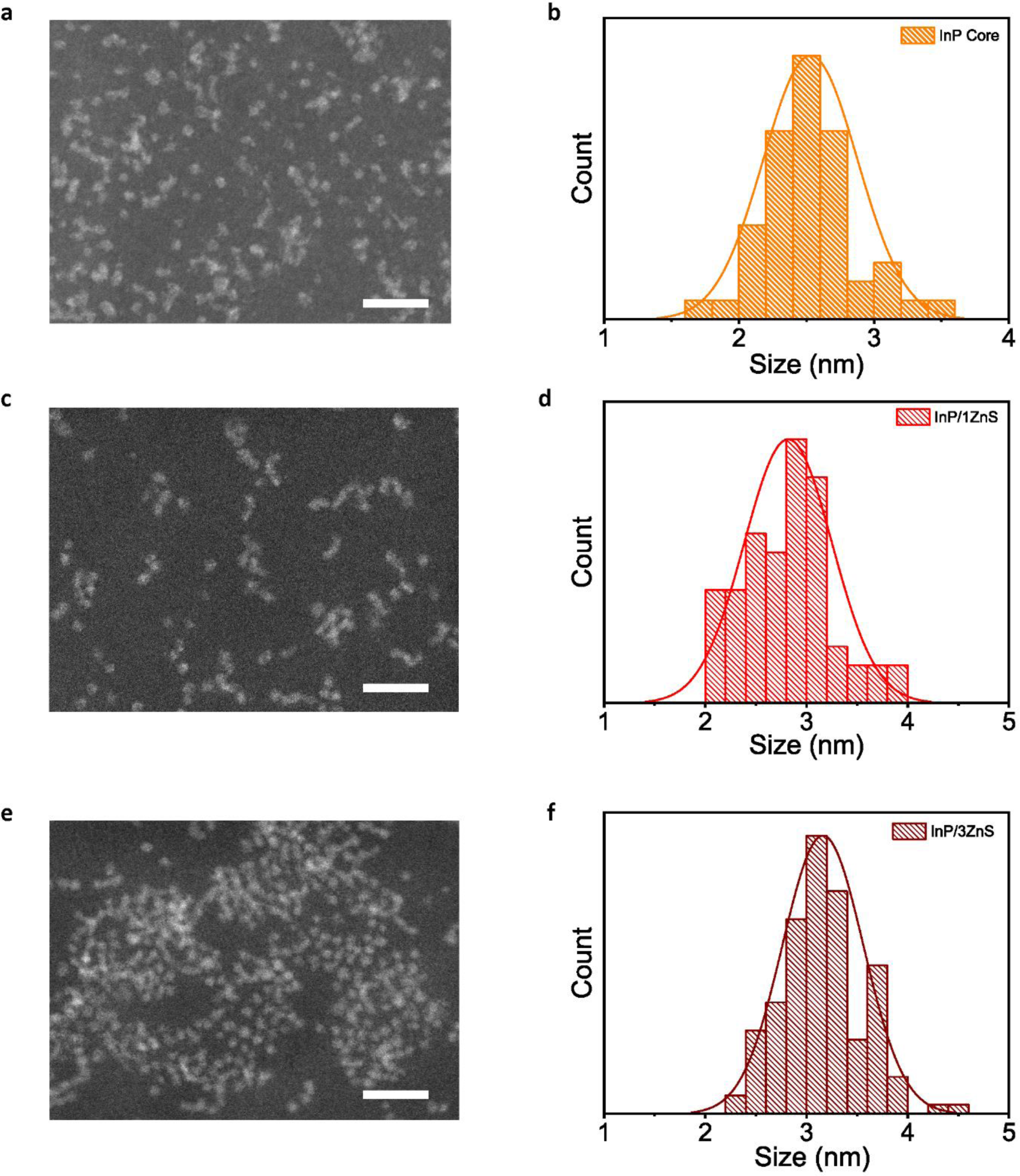
**(a)** TEM image of InP core QDs. **(b)** Size distribution of InP core QDs where the mean size is 2.49 nm. **(c)** TEM image of InP/0.5ZnS core/shell QDs. **(d)** Size distribution of InP/0.5ZnS core/shell QDs where the mean size is 2.81 nm. **(e)** TEM image of InP/1ZnS core/shell QDs. **(f)** Size distribution of InP/3ZnS core/shell QDs where the mean size is 3.15 nm. 200 QDs are counted for each size distribution plot. Scale bars of TEM images are 20 nm.

**Figure S3.**
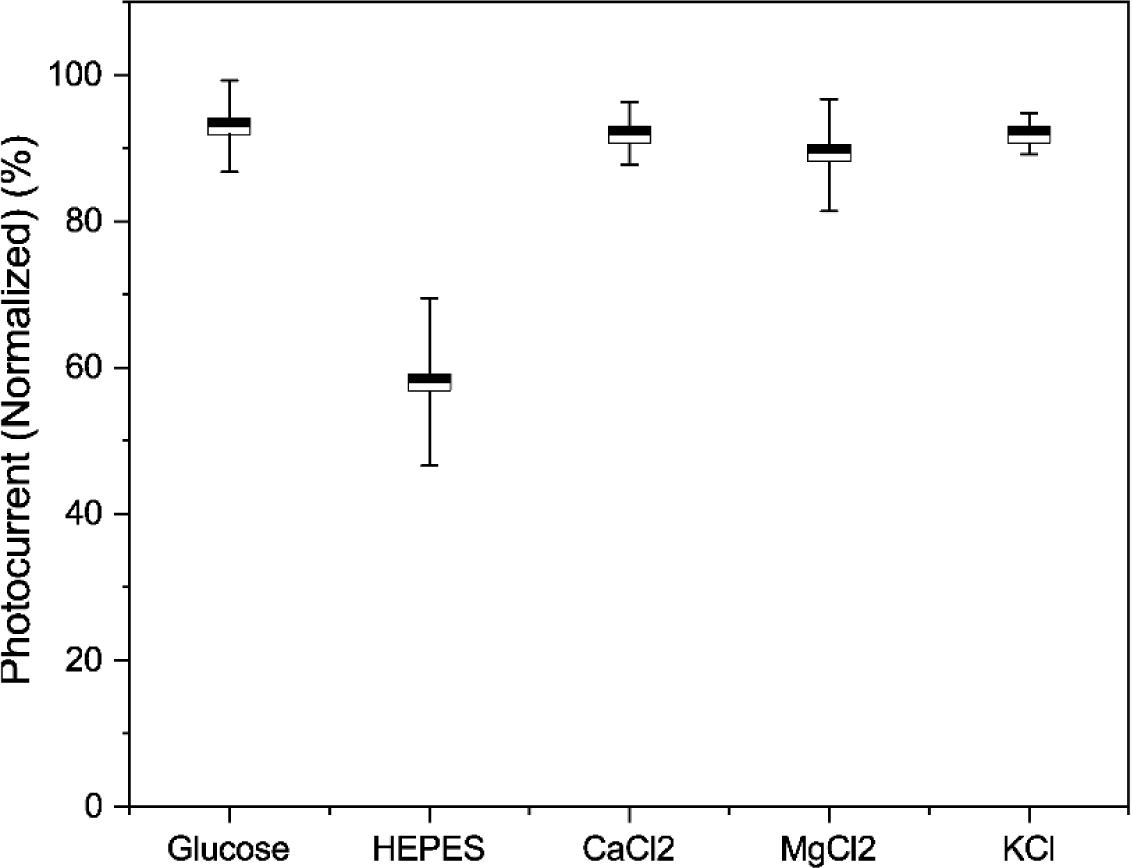
Photocurrents while each ingredient is decreased to their half (means ± SD, n=5)

**Figure S4.**
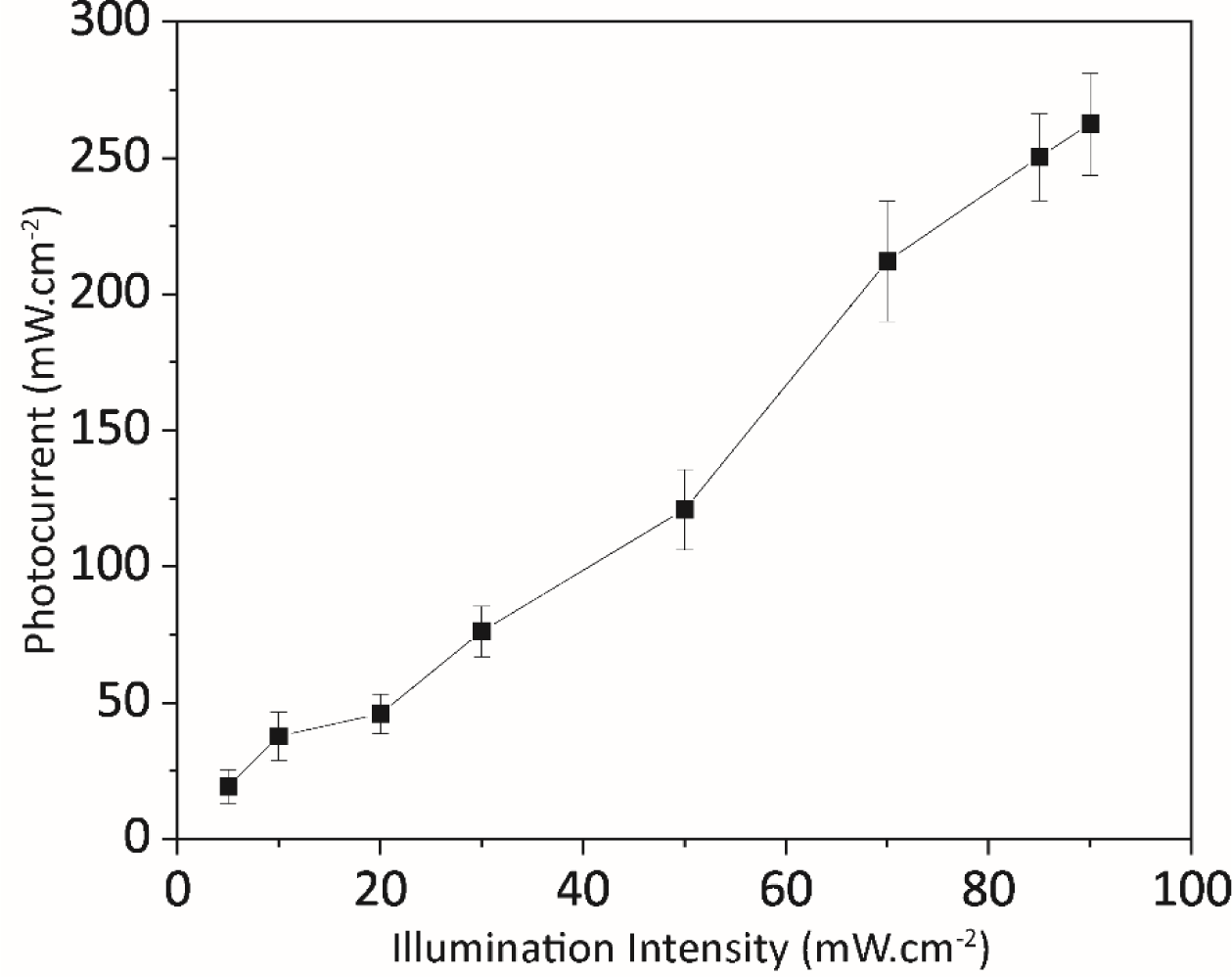
Photocurrent measurements in the aCSF electrolyte under 20 ms illumination under different light intensities (means ± SD, n=5)

**Figure S5.**
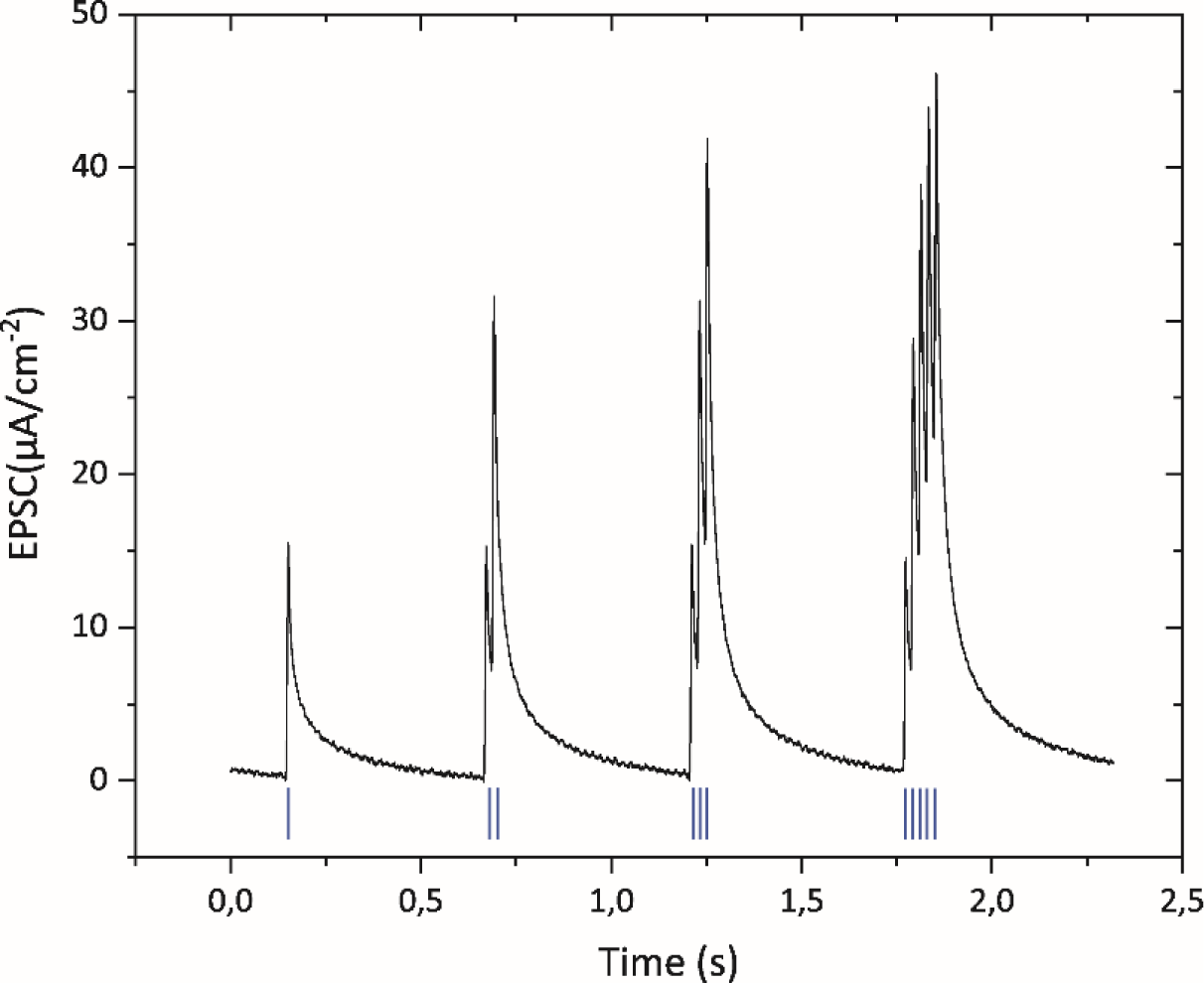
Spike-number-depended-plasticity (SNDP) for 1,2,3, and 5 pulses.

**Figure S6.**
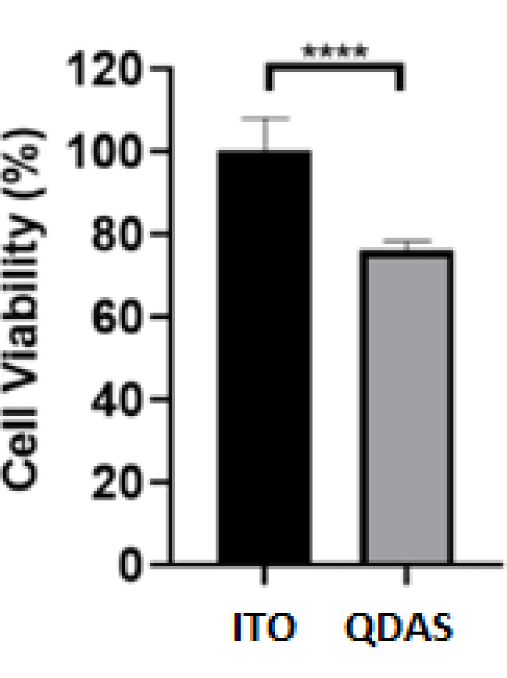
Biocompatibility analysis of QDAP compared to ITO control. CTG cell viability result of primary hippocampal neurons cultured on QDAPs and ITO substrates (mean ± SD for n = 4). An unpaired, two-tailed t-test was used for statistical analysis, and ****p < 0.0001 was evaluated as statistically significant. ∼80% biocompatibility indicates that it is generally well-tolerated by cells and has a relatively low level of adverse effects. Such biocompatibility levels are appropriate to grow primary neurons on the device for photostimulation [1].

**Figure S7.**
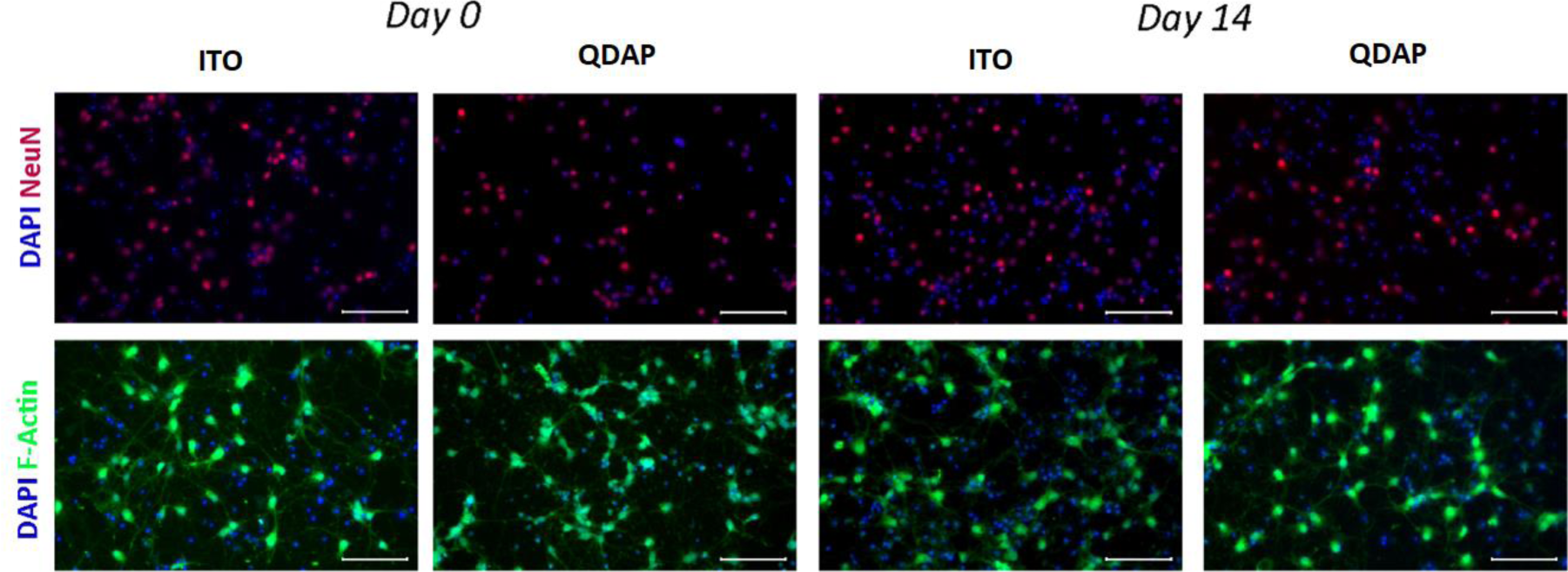
Immunofluorescence images of primary hippocampal neurons cultured on QDAP and ITO substrates on days 0 and 14 of culturing to observe morphology and viability of primary hippocampal neurons. Cells were co-stained with DAPI (blue) to show the nucleus, Anti-NeuN (red) to show the neuronal nucleus, and Anti-f-Actin (green) to indicate cell structure (scale bar: 100 μm).

**Figure S8.**
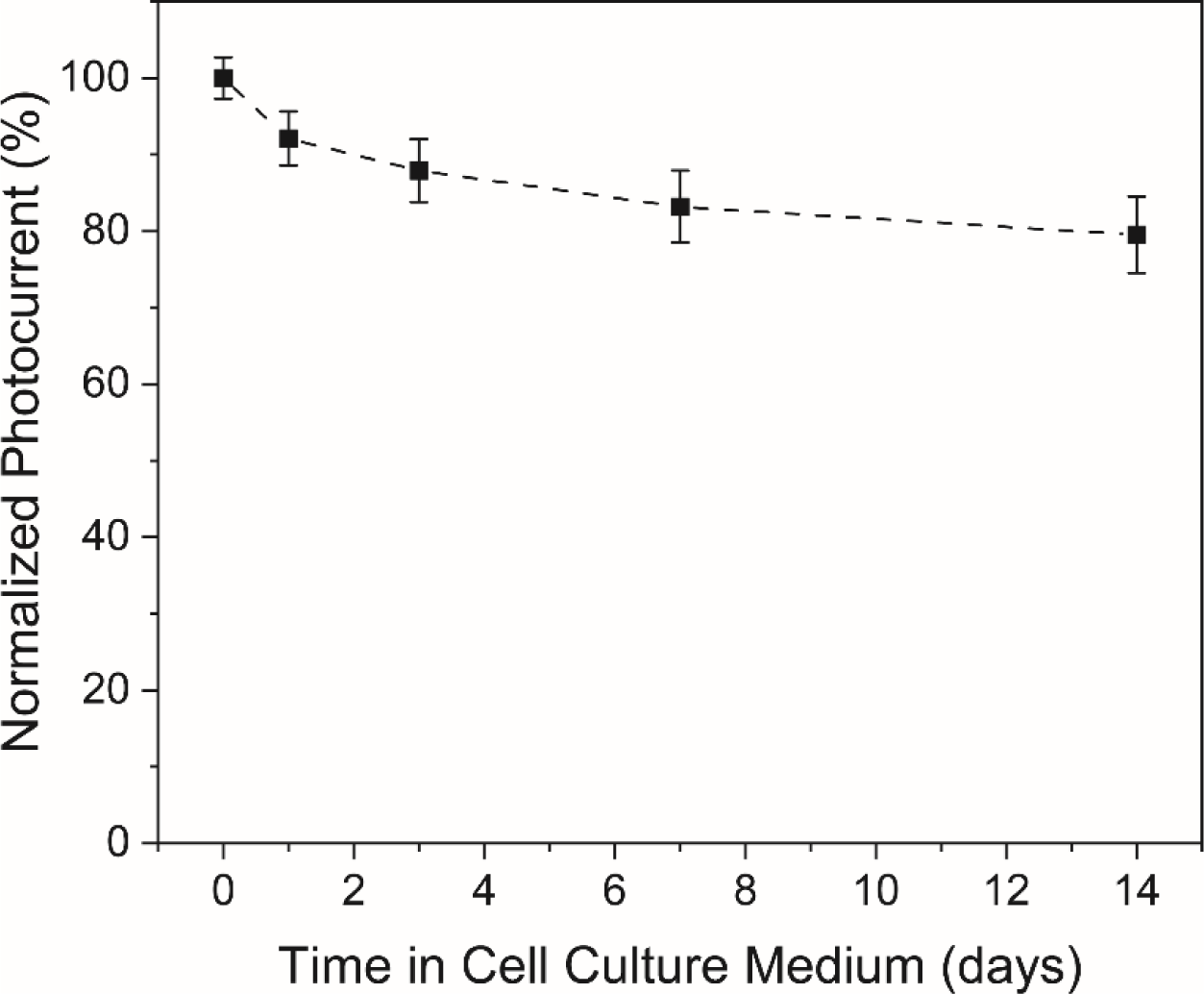
Normalized Photocurrent peak up to 14 days of cell culture.

**Figure S9.**
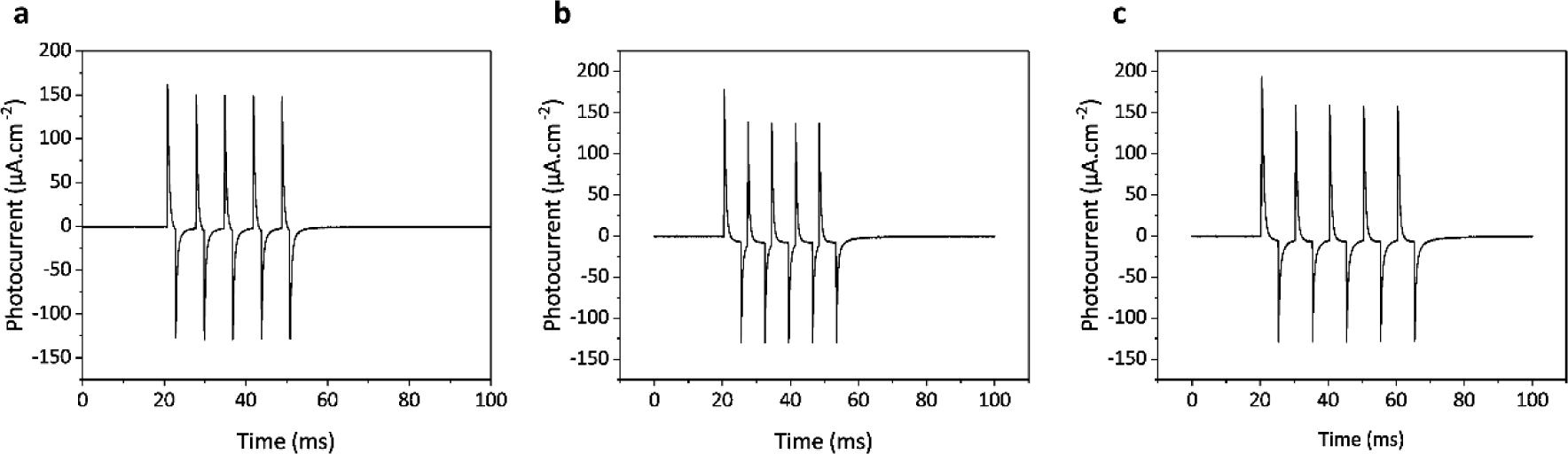
Capacitive photocurrent response of ITO/ZnO/P3HT device triggered with different illuminations, (left) illuminated with 2 ms on time and 5 ms off time, (middle) illuminated with 5 ms on time and 5 ms off time, (right) illuminated with 5 ms on time and 2 ms off time.

## Notes

### Competing Interest Statement

The authors have declared no competing interest.

