## Supplementary material for "A Retina-inspired Optoelectronic Synapse Using Quantum Dots for Neuromorphic Photostimulation of Neurons": Supplemetary info

**This PDF file includes:**

Supplementary Text

Figs. S1 to S9

Tables S1

References (1)

**Table S1.** Zinc and sulfur precursor solutions used for ZnS shelling.

| | **Concentration (M)** | **InP/0.5ZnS** | **InP/1ZnS** |
|---|---|---|---|
| **$ZnSt_2$-ODE** | 0.1 | 600 µL | 1050 µL |
| **S-TOP** | 0.1 | 600 µL | 1050 µL |

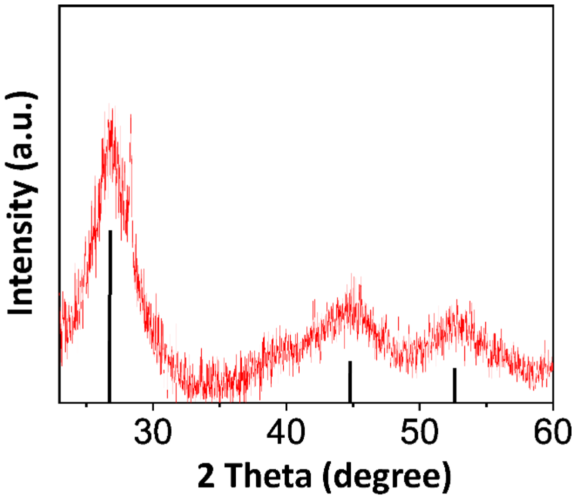


**Figure S1.** XRD pattern of the InP/ZnS core/shell QDs. The X-ray diffraction (XRD) pattern reveals the crystal planes of the (111), (220), and (222) of the QDs. (InP JCPDS No. 32-0452 and ZnS JCPDS No. 80-0020)

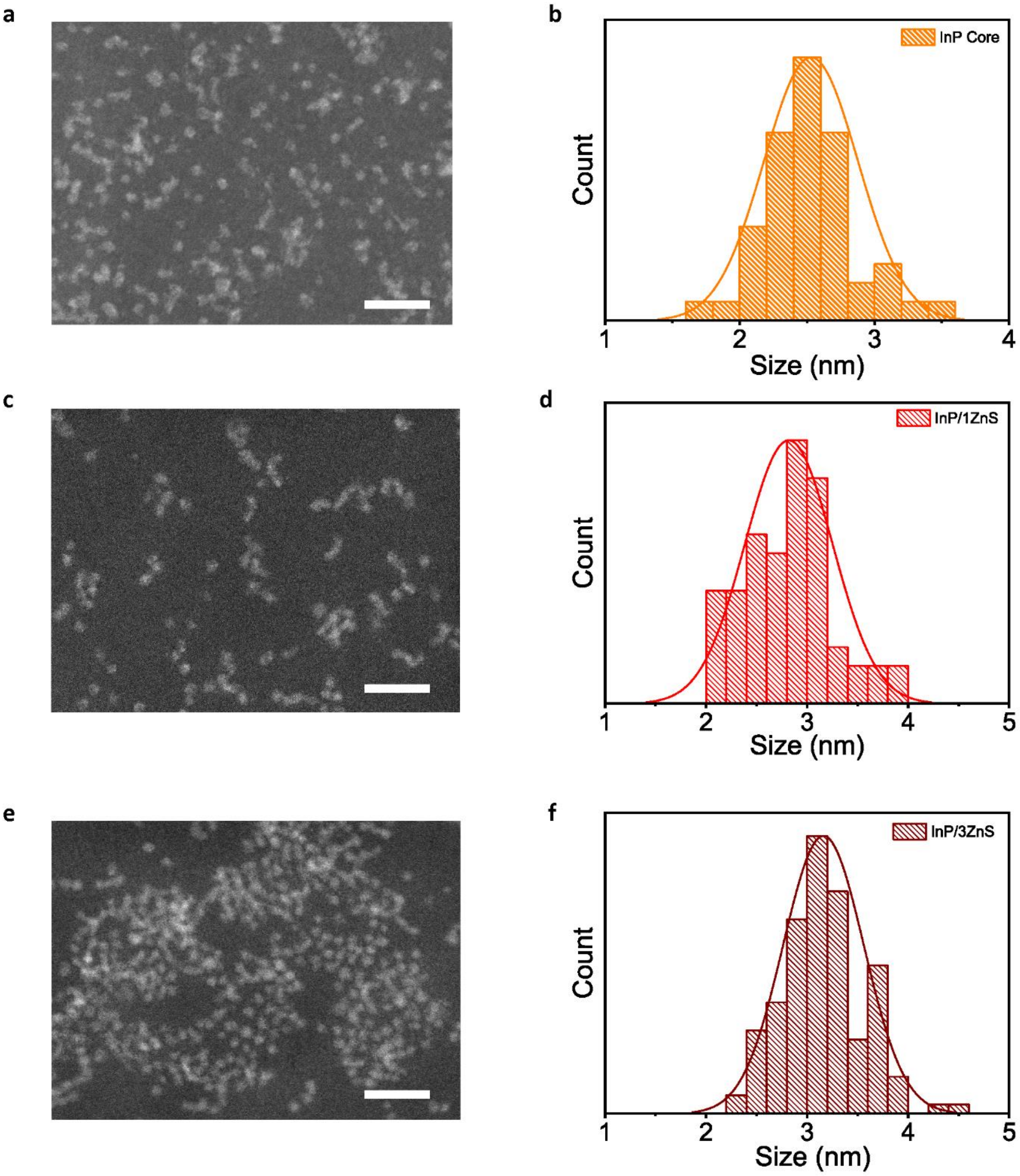


**Figure S2. (a)** TEM image of InP core QDs. **(b)** Size distribution of InP core QDs where the mean size is 2.49 nm. **(c)** TEM image of InP/0.5ZnS core/shell QDs. **(d)** Size distribution of InP/0.5ZnS core/shell QDs where the mean size is 2.81 nm. **(e)** TEM image of InP/1ZnS core/shell QDs. **(f)** Size distribution of InP/3ZnS core/shell QDs where the mean size is 3.15 nm. 200 QDs are counted for each size distribution plot. Scale bars of TEM images are 20 nm.

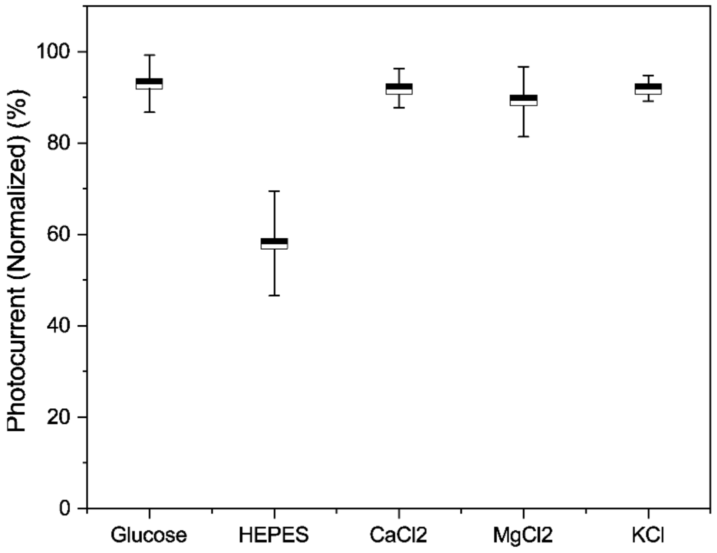


**Figure S3.** Photocurrents while each ingredient is decreased to their half (means ± SD, n=5).

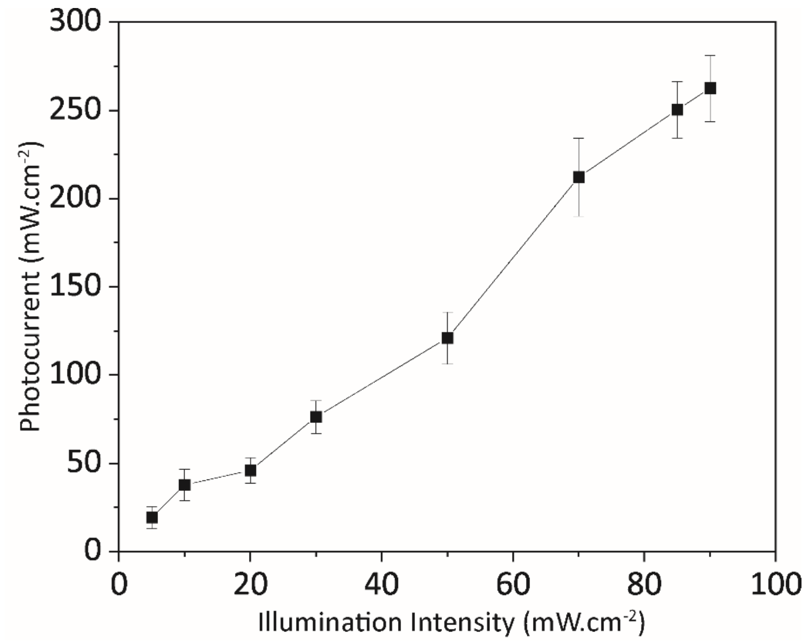


**Figure S4.** Photocurrent measurements in the aCSF electrolyte under 20 ms illumination under different light intensities (means ± SD, n=5).

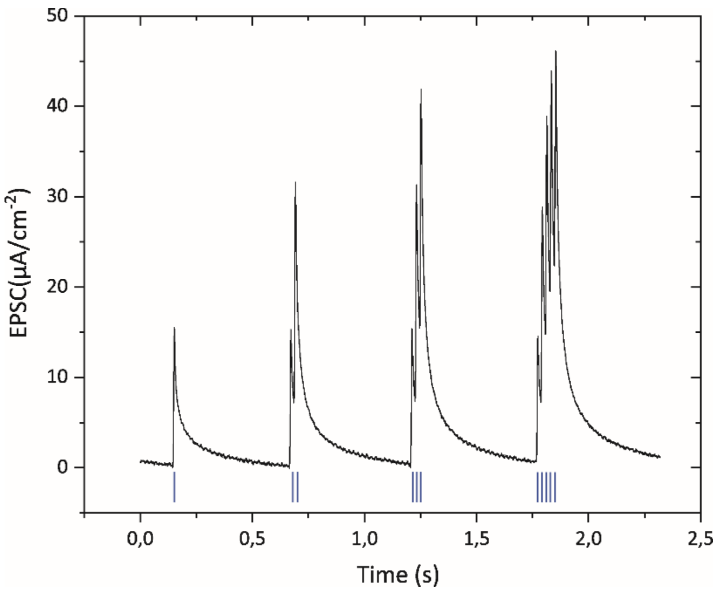


**Figure S5.** Spike-number-depended-plasticity (SNDP) for 1,2,3, and 5 pulses.

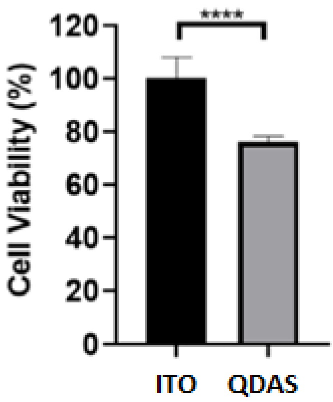


**Figure S6.** Biocompatibility analysis of QDAP compared to ITO control. CTG cell viability result of primary hippocampal neurons cultured on QDAPs and ITO substrates (mean ± SD for n = 4). An unpaired, two-tailed t-test was used for statistical analysis, and ****$p < 0.0001$ was evaluated as statistically significant.

~80% biocompatibility indicates that it is generally well-tolerated by cells and has a relatively low level of adverse effects. Such biocompatibility levels are appropriate to grow primary neurons on the device for photostimulation [1].

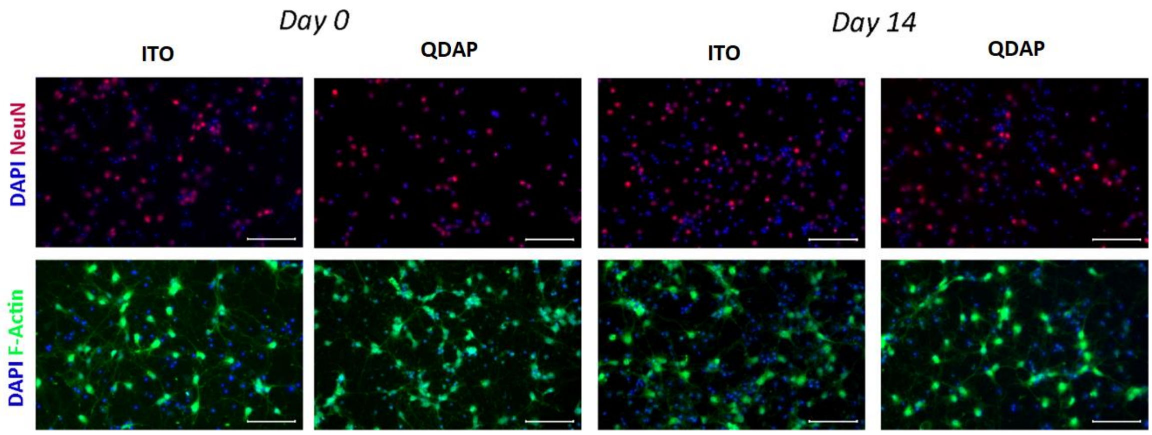


**Figure S7.** Immunofluorescence images of primary hippocampal neurons cultured on QDAP and ITO substrates on days 0 and 14 of culturing to observe morphology and viability of primary hippocampal neurons. Cells were co-stained with DAPI (blue) to show the nucleus, Anti-NeuN (red) to show the neuronal nucleus, and Anti-f-Actin (green) to indicate cell structure (scale bar: 100 μm).

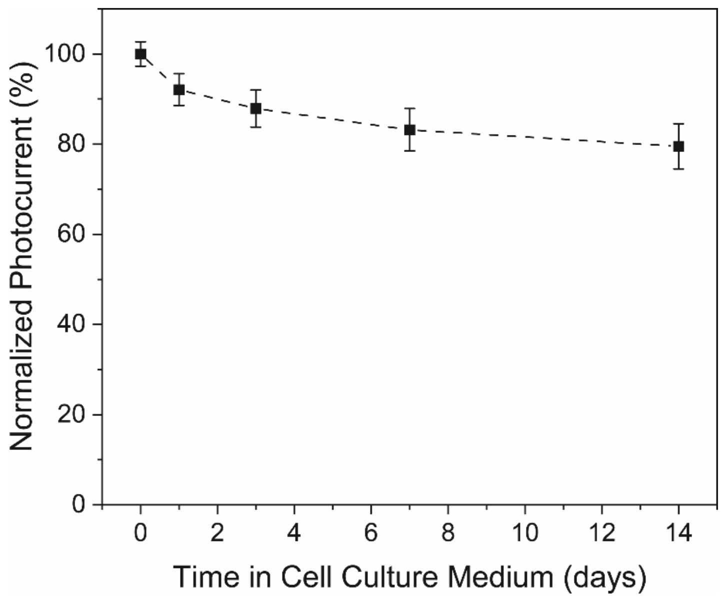


**Figure S8.** Normalized Photocurrent peak up to 14 days of cell culture.

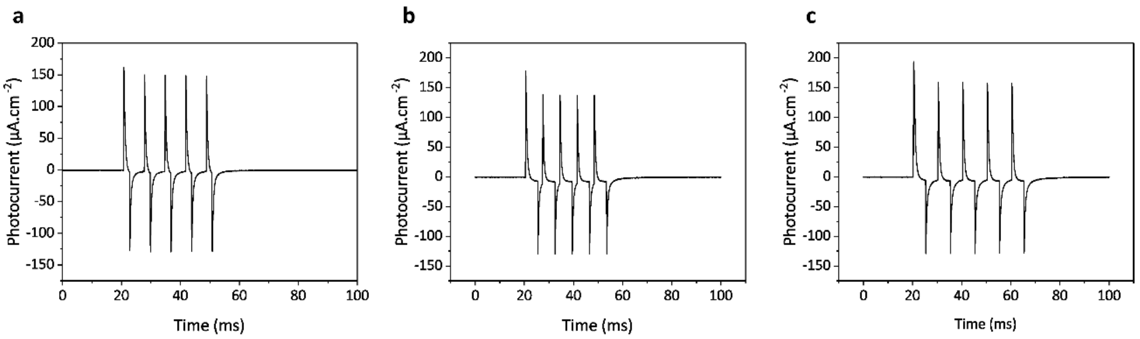


**Figure S9.** Capacitive photocurrent response of ITO/ZnO/P3HT device triggered with different illuminations, (left) illuminated with 2 ms on time and 5 ms off time, (middle) illuminated with 5 ms on time and 5 ms off time, (right) illuminated with 5 ms on time and 2 ms off time.
